# Age influences audiovisual speech processing in multi-talker scenarios – Evidence from cortical oscillations

**DOI:** 10.1101/2022.02.23.481314

**Authors:** Alexandra Begau, Laura-Isabelle Klatt, Daniel Schneider, Edmund Wascher, Stephan Getzmann

## Abstract

Age-related differences in the processing of audiovisual speech in a multi-talker environment were investigated analyzing event-related spectral perturbations (ERSPs), focusing on theta, alpha, and beta oscillations that are assumed to reflect conflict processing, multisensory integration, and attentional mechanisms, respectively. Eighteen older and 21 younger healthy adults completed a two-alternative forced-choice word discrimination task, responding to audiovisual speech stimuli. In a cocktail-party scenario with two competing talkers (located at-15° and 15° azimuth), target words (/yes/ or /no/) appeared at a pre-defined (attended) position, distractor words at the other position. In two audiovisual conditions, acoustic speech was combined either with congruent or uninformative visual speech. While a behavioral benefit for congruent audiovisual speech occurred for both age groups, differences between audiovisual conditions in the theta and beta band were only present for older adults. A stronger increase in theta perturbations for stimuli containing uninformative visual speech could be associated with early conflict processing, while a stronger suppression in beta perturbations for congruent audiovisual speech could be associated to audiovisual integration. Compared to the younger group, the older group showed generally stronger beta perturbations. No condition differences in the alpha band were found. Overall, the findings suggest age-related differences in audiovisual speech integration in a multi-talker environment. While the behavioral benefit of congruent audiovisual speech was unaffected by age, older adults had a stronger need for cognitive control when processing conflicting audiovisual speech input. Furthermore, mechanisms of audiovisual integration are differently activated depending on the informational content of the visual information.

## 1 Introduction

Audiovisual speech scenarios with multiple talkers occur to us frequently in our daily lives – whether it is during a meeting with friends for dinner or in highly digitalized working and social environments, where audiovisual online meetings and conferences become the default. When listening to a person in a multi-talker situation, our cognitive system needs to select the relevant speech stream and to focus attention on this speech information, while suppressing all the irrelevant streams and back-ground noise (Bregman & McAdams, 1994; Bronkhorst, 2015). Under these “cocktail-party” conditions (Cherry, 1953), audiovisual speech may provide a benefit in speech comprehension against unimodal speech which appears to be even more pronounced in hearing-impaired and older people (Begau et al., 2021; van Wassenhove et al., 2005; Winneke & Phillips, 2011). Thus, despite age-related sensory and cognitive decline in speech perception, older adults’ ability to integrate multisensory linguistic information usually remains intact and can be used as a compensatory mechanism (Friedman, 2011; Passow et al., 2012; Winneke & Phillips, 2011). The mechanisms of audiovisual speech processing are closely intertwined with neural oscillatory brain activity, such as theta, alpha and beta oscillations (Keil & Senkowski, 2018). In the following study, we employed an audiovisual multi-talker scenario with young and old participants, varying the informational content of the visual or auditory modality, and investigated age differences and an audiovisual benefit indicated in event-related spectral perturbations (ERSPs).

### 1.1 Audiovisual multi-talker scenarios and the role of age

Naturally, speech occurs multimodally, providing both auditory and visual information. The integration of audiovisual speech requires several processing steps, including feed forward -feedback loops, attentional modulation and predictive coding (see the framework by Keil & Senkowski, 2018). Visual speech typically precedes auditory speech, especially at the very beginning of a verbal utterance (Chandrasekaran et al., 2009; Schwartz & Savariaux, 2014), adding to the redundancy of the signal and enabling the formation of predictions (Bronkhorst, 2015; Peelle & Sommers, 2015). In particular, preceding visual information enables *predictive coding* (Arnal & Giraud, 2012), forming expectations on the anticipated auditory speech information. This is in line with the *analysis by synthesis* hypothesis by Wassenhove and colleagues (2005)suggesting an early integration process. The more redundant and thus predictive the value of the visual information is, the higher the expected audiovisual benefit. The neuro-cognitive mechanisms involved in processing and integrating audiovisual speech in multi-talker scenarios are susceptible to age-related changes. Aging in general is associated with a decline in sensory (for review, audition: Martin & Jerger, 2005; vision: Owsley, 2011), behavioral and cognitive abilities (see Friedman, 2011). While pre-attentive filtering in multi-talker scenarios is usually not affected in healthy aging, age-related deficits become apparent in higher-order attentional functions, such as the top-down modulation of information (Peelle & Wingfield, 2016). Several studies found age-related deficits when maintaining and coordinating competing information, processing irrelevant information and focusing attention on relevant stimuli (Correa-Jaraba et al., 2016; see review by Guerreiro et al., 2010; Rey-Mermet & Gade, 2018). As a result, older adults experience higher conflict costs and more difficulties in tasks with high attentional demand (Passow et al., 2012). However, despite these age-related difficulties in speech comprehension (Getzmann et al., 2020; Martin & Jerger, 2005), audiovisual integration per se appears to be still intact, offering a possibility to compensate for age-related decline (Begau et al., 2021; Cienkowski & Carney, 2002; Sekiyama et al., 2014; Sommers et al., 2005; Winneke & Phillips, 2011; Wong et al., 2010). In fact, there are studies indicating that older adults make use of additional visual information to an even greater degree than young adults (Sekiyama et al., 2014), especially when auditory information is ambiguous (Cienkowski & Carney, 2002).

### 1.2 Neural oscillations as correlates of audiovisual speech processing

The timing of the different steps of multisensory speech processing and integration have been mostly studied using event-related potentials (Baart et al., 2014; Begau et al., 2021; Stekelenburg & Vroomen, 2007; van Wassenhove et al., 2005; Winneke & Phillips, 2011). Recently, oscillatory activity in the brain has gained more attention in relation to multisensory integration, possibly reflecting the communication between different cortical areas during processing (for review see Keil & Senkowski, 2018). For example, in an audiovisual cocktail-party setting, the involvement of oscillatory brain activity with frequencies between 0.5 and 15 Hz was shown, suggesting that viewing the speakers face enhances the capacity of the auditory cortex to selectively represent and track the relevant speaker (Zion Golumbic et al., 2013). For the present study, we focused on the role of theta, alpha and beta oscillations for audiovisual processing. In the following sections, we briefly summarize the functions of neural oscillations in the processing of audiovisual speech stimuli.

#### 1.2.1 Theta

Midfrontal theta (4-8 Hz) is thought to reflect communication between different brain areas when the need for cognitive control arises (Cavanagh & Frank, 2014). An increase in theta power can be observed in tasks with high working memory load (Gevins, 1997; Jensen & Tesche, 2002; Maurer et al., 2015; for audiovisual speech stimuli Michail et al., 2021). In audiovisual tasks, theta band activity has been attributed to divided attention (Keller et al., 2017) and the encoding of new information and change detection (Fingelkurts et al., 2007). In illusory McGurk stimuli, where non-congruent auditory and visual information such as /ba/ and /ga/ form a merged, illusory percept /da/ (McGurk & MacDonaldJohn, 1976) but also with incongruent audiovisual information, an increase in midfrontal theta can be observed, supporting the notion of a conflict processing network (Keil et al., 2012; Morís Fernández et al., 2018). This increase in theta power is further interpreted as a feedback mechanism, which comes into play when incongruent information is not simply ignored (Lange et al., 2013), but is integrated and modulates the audiovisual percept (Morís Fernández et al., 2015). In older adults, midfrontal theta is decreased during retention and recognition, suggesting theta power to be a relevant marker for task-specific cognitive aging (Cummins & Finnigan, 2007). To summarize, midfrontal theta increase is associated with a need for cognitive control, followed by top-down attentional modulation of perception to enable successful audiovisual integration (Friese et al., 2016; Keil & Senkowski, 2018).

#### 1.2.2 Alpha

Higher cognitive load is usually not only associated with an increase in theta power, but also a suppression in alpha power (8 – 12 Hz; e.g., Wascher et al., 2019). This complementary occurrence of alpha and theta can be observed in conflict processing, with theta being associated with the processing of a conflict or a mismatch, and alpha being associated with the maintenance of the stimulus representation(Gratton, 2018). Furthermore, Keller and colleagues (2017) discussed the role of alpha and theta oscillations in multisensory attention. While frontocentral theta represents divided attention, which is necessary for the simultaneous processing of both auditory and visual input, alpha oscillations – especially over parieto-occipital scalp sites – seem to play an important role in selective attention, such as the focusing on relevant and suppression of irrelevant auditory information (Klatt et al., 2020; Schneider et al., 2021; Wöstmann et al., 2016). Specifically, a posterior alpha power increase can be found when irrelevant working memory contents are suppressed (Klatt et al., 2020). Increased probability for alpha oscillations occurring with the presentation of audiovisual stimuli relative to auditory-only stimuli is interpreted as a higher need for focused attention in the maintenance of audiovisual stimuli (Fingelkurts et al., 2007). Meanwhile, a decrease in parietal alpha has been found in speech stimuli with increasing predictiveness and decreasing acoustic detail (Wöstmann, Herrmann, Wilsch, & Obleser, 2015). Weaker alpha suppression can further be found in degraded (Weisz et al., 2011) or less intelligible speech (Drijvers et al., 2018). Moreover, stronger medio-central alpha suppression can be found in McGurk illusion trials compared to congruent audiovisual stimuli (Roa Romero et al., 2016). In older participants, delayed alpha activity in multi-talker speech processing has been linked to cognitive slowing (Getzmann et al., 2020), and weaker alpha power modulations have been associated with difficulties listening to speech in noise, with older listeners being more affected by variations in acoustic detail (Wostmann et al., 2015). Taken together, posterior alpha oscillations are likely associated with the successful selection of the targeted speech and the focus of attention on relevant information.

#### 1.2.3 Beta

Oscillations in the beta band (16 to 30 Hz) have been shown to be closely intertwined with alpha oscillations during tasks requiring attentional inhibition in multisensory tasks (Friese et al., 2016; Ganesan et al., 2021). Beta oscillations are associated with neural networks responsible for error-monitoring and decision making (Arnal & Giraud, 2012; Friese et al., 2016) and the maintenance of current sensorimotor information (Engel & Fries, 2010; Schneider et al., 2020). A decrease in beta power can be observed following stimulus-driven stimulation (Engel & Fries, 2010), and is stronger in audiovisual conditions with background noise (Schepers et al., 2013) as well as following McGurk illusions (Roa Romero et al., 2015). In audiovisual speech stimuli using McGurk illusions, an early and a late beta power suppression can be distinguished (Michail et al., 2021; Roa Romero et al., 2015). While the early suppression is interpreted as the formation of a fusion percept in McGurk illusions and error processing in incongruent audiovisual stimuli, late beta suppression is interpreted as a marker for top-down audiovisual integration (Michail et al., 2021). Beta oscillations have been associated with networks related to executive function, with a decrease in power in older adults, together with a posterior-to-anterior shift of activation (Enriquez-Geppert & Barceló, 2018). Taken together, beta band oscillations appear to play a role in monitoring information to form an integrated audiovisual percept (Keil & Senkowski, 2018).

### 1.3 Research Question and Hypotheses

So far, very little is known about how our brain processes audio-visual speech especially in multi-talker situations and whether age plays a role for the integration of audio-visual speech. In the present study, we investigated multimodal speech processing in a simulated cocktail-party situation, in which younger and older participants responded to short speech stimuli in a speeded two-alternative forced-choice word discrimination paradigm. We contrasted two audiovisual conditions, in which the visual information was either congruent (i.e., contained the same information as the auditory information), or did not contain relevant information. We focused on oscillatory correlates of audiovisual speech processing in theta, alpha, and beta bands and were especially interested in age-related differences. For behavioral measures, we expected better performance for audiovisually congruent compared to uninformative additional visual input, that is, faster responses and higher accuracy. Even though we expected a general age-related decline in performance, older adults should profit more strongly from audiovisually congruent information than young adults. For oscillatory modulations, we focused on event-related spectral perturbations (ERSPs), that is “mean change in spectral power (in dB) from baseline” (Makeig et al., 2004). In line with previous work (e.g. Cavanagh & Frank, 2014; Keil et al., 2012; Morís Fernández et al., 2018), we expect an increase in theta perturbations over frontocentral areas after sound onset. Since frontocentral theta has been associated with conflict processing and the need for cognitive control, we expect this increase to be larger for uninformative visual compared to congruent audiovisual stimuli. In line with the above-mentioned work (e.g. Keller et al., 2017; Misselhorn et al., 2019), audiovisual facilitation should also be reflected in frontal and parieto-occipital alpha band activity: Assuming that congruent audiovisual information requires less attentional resource allocation for successful processing of the stimulus, we expect ERSPs to show less alpha power suppression for audiovisually congruent stimuli compared to uninformative visual information. Similarly (in accordance with e.g. Roa Romero et al., 2015), we expect ERSPs to show a beta power suppression after sound onset, which should be less pronounced for uninformative visual compared to congruent audiovisual stimuli. Furthermore, we expect a general aging effect to be reflected in ERSPs indicating a weaker increase in theta power and alpha and beta power suppression in older compared to younger adults. Finally, assuming older participants profit more from congruent audiovisual information than younger adults, we expect a larger difference between the audiovisual conditions in the older than in the younger group.

## 2 Methods

### 2.1 Participants

Out of 50 invited participants, eleven were excluded from the analysis due to technical issues (*n* = 2), hearing (*n* = 4) and visual problems (*n* = 1), insufficient compliance to complete the experiment (*n* = 1), misunderstood instructions (*n* = 2) and responses below chance level (*n* = 1). In total, *N* = 39 participants were analyzed. Participants in the older group (*n* = 18), were 60 to 70 years old (*M* = 65.39, *SE* = 0.85; 9 male), the younger group (*n* = 21) was aged 20 to 34 years (*M* = 25.29, *SE* = 0.72; 9 male). All participants reported to be right-handed and without neurological or psychiatric illness. Prior to testing, written informed consent was given by each participant. Written informed consent was given prior to testing, the experiment was approved by the Ethical Committee of the Leibniz Research Centre for Working Environment and Human Factors, Dortmund, Germany and in accordance with the Declaration of Helsinki. 10€ per hour were paid at the end of the experiment.

### 2.2 Sensory and cognitive abilities

To assess the participants’ hearing level, each participant completed a pure-tone audiometry (Oscilla USB 330; Immedico, Lystrup, Denmark) with eleven pure-tone frequencies from 125 to 8000 Hz. Hearing thresholds were mostly ≤ 30dB for frequencies below 4000 Hz, with mild to moderate presbycusis in the older group. Overall, we observed hearing levels of ≤ 35dB at 2000 Hz (n = 2) and ≤ 35 dB (*n* = 2), ≤ 40 dB (*n* = 2) and ≤ 45 dB (*n* = 1) at 3000 Hz. According to WHO criteria (Olusanya et al., 2019) we evaluated the hearing threshold averaged across 500, 1000, 2000 and 4000 Hz in the better ear, demonstrating that hearing can be considered as unimpaired in both the younger group (mean 7.37 dB, *range* = 2.5 – 12.5, *SE* = 0.70) and in the older group (mean threshold 15dB, *range* = 3.75 – 21.25, *SE* = 1.11). However, not surprisingly, the older participants had significantly higher hearing thresholds than the younger ones, *t*(37) = 5.85, *p* < .001, *g* = 1.83. (Olusanya et al., 2019). Due to the relatively loud stimulus presentation at 75dB, outliers in hearing level for individual frequencies are considered acceptable.

Landolt C optotypes at 1.5m distance were used to measure visual acuity. All but one (older) participant reached a value of 1.0 or above, which is considered normal acuity (ISO 8596:2017(E)). Visual acuity was significantly better in the younger compared to the older group. Additionally, we assessed contrast sensitivity using a *Pelli-Robson Contrast Sensitivity Chart* in 1.0m distance (Pelli, Robson, & Wilkins, 1988). All but one (older) participant scored the recommended contrast sensitivity of 1.65 or above (Mäntyjärvi & Laitinen, 2001). Younger participants showed better visual acuity, *V* = 308.5, *p* <.001, *r* = .56 and contrast sensitivity, *V* = 327, *p* <.001, *r* = .68, than older participants.

To assess cognitive function, we chose the *Montreal Cognitive Assessment* (MoCA; Nasreddine et al., 2005), a screening for mild cognitive impairment. In the older group, scores between 19 and 30 (*M* = 26.39, *SE* = 6.22) were reached. Younger participants performed significantly better with scores between 26 and 30 points (*M* = 28.76, *SE* = 6.28, *V* = 77.5, *p* = .001, *r* = .51). Overall, there was no indication of dementia (i.e., a sum score of 17 points or lower Carson et al., 2018).

### 2.3 Materials and stimuli

The experiment took place in a sound attenuated room (5.0 × 3.3 × 2.4 m^3^) with a background noise level of 20 dB (A). Ceiling and walls were covered in pyramid shaped foam panels and a woolen carpet to achieve attenuation. Participants were seated in front of a horizontal array consisted of two 12” vertically aligned monitors (1080 × 1920 px, 50Hz refresh rate; Beetronics, Düsseldorf, Germany) on a left and right position (−15° and +15° azimuth, fig.1A). The auditory signal was displayed at a level of 75dB SPL from full range loudspeakers; one mounted under each monitor (SC 55.9 – 8 Ohm; Visaton Haan, Germany). The setup was in 1.5m distance at head height (1.00 m (loudspeaker) and 1.12 m (monitor)) from the participant’s seat.

**Figure 1:**
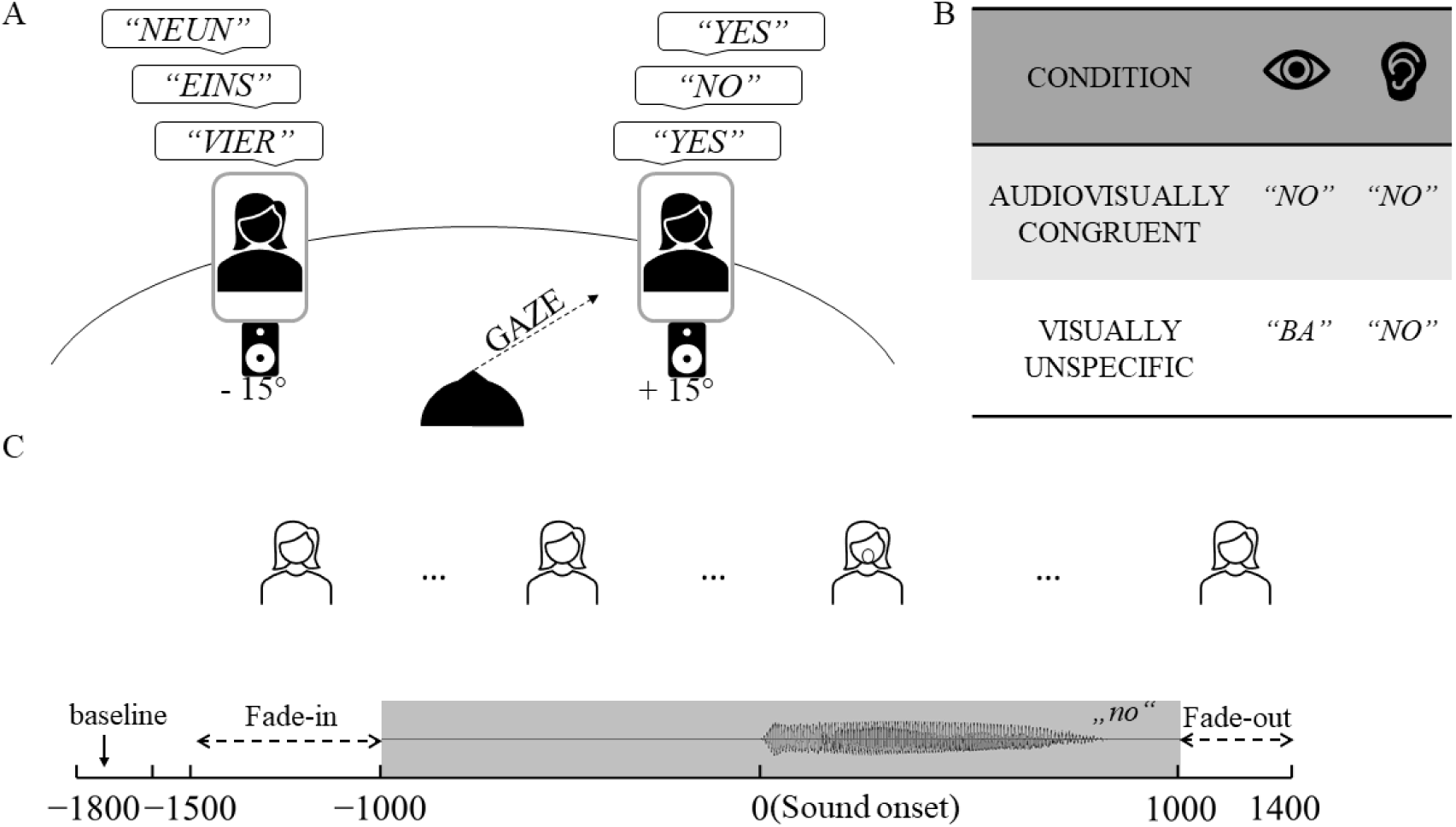
Experimental design and audiovisual stimuli: (A) Laboratory setup with brief target words (/yes/ and /no/) presented at a pre-defined location (here +15°), and distracting words (German number words) at the opposite position. (B) Audiovisual stimulus conditions and manipulation of visual speech, being either congruent to acoustic speech (here /no/) or non-informative (/ba/). (C) Structure of (epoched) trials, time-locked to the onset of acoustic speech.

In the same room, close-ups of two female speakers showing the face and neck of the respective speaker were recorded in front of a grey-blue background (RGB 106, 145, 161, HDR-CX220, Sony, Tokyo, Japan; 1920 × 1080-pixel resolution, 50fps frame rate). The audio was recorded simultaneously using a dynamic USB-microphone (Podcaster, RØDE, Silverwater, NSW, Australia; mono, 48kHz, 24-bit). Both speakers were German dialect-free natives with a fundamental frequency of 169 Hz (*SE* = 2.00) and 213 Hz (*SE* = 7.87), respectively. The stimulus material consisted of the speakers uttering short words; /yes/ and /no/ (in English) were assigned target words and German digit words from /one/ to /ten/ were distractor words. The speakers were instructed on clear accentuated pronunciation with a neutral facial expression and no head movements. Target words were chosen due to their clear difference in articulation both auditory and visual thus being well distinguishable. The auditory signal was recorded and edited in Audacity (version 2.3.0), applying noise reduction (15 dB) and normalization (−6 dB). Audio and video tracks were then merged and further edited in Shotcut (libx264/aac codec, export as MP4). For the *audiovisually congruent condition*, videos were cut from the on-to offset of visible lip movement, with lips being closed at the start and end of a stimulus video. 500 ms of fade-in and 400 ms of fade-out from and to black were added to the tails of the video, using the first and last video frame, respectively. Additionally, we added a variable amount of freeze frames between fades and speech video, resulting in a total video length of 2900 ms with the auditory onset being locked to 1500 ms. Thus, a synchronous onset of auditory speech was ensured during the experiment. In the final stimulus material (fig. 1C), the auditory speech had an average length of 555.33 ms (*range* = 486-702 ms, *SE* = 9.93), while the visual speech was at average 1120.91 ms long (*range* = 740 -1540 ms, *SE* = 53.76). The onset of lip movement preceded auditory speech on average for 430.91 ms (*range* = 120-860 ms, *SE* = 44.40 ms). To manipulate the audiovisual speech information for the *visually unspecific* condition, we further edited the stimulus material, creating stimuli with incongruent information (fig. 1B). Here, we replaced the speech video and the variable freeze frames with a video of the respective speaker saying /ba/ (lip movement: 1360 and 600 ms, latency between lip and sound onset: 660 and 160 ms, respectively), resulting in distinguishable auditory information, but identical visual information for all stimuli.

### 2.4 Experimental Procedure

Prior to testing, hearing level, visual acuity, contrast sensitivity and cognitive abilities were assessed. Participants were then prepared for the EEG recording and received both written and oral instructions about the task. Participants were told to respond as fast and as accurate as possible, operating a two-alternative forced choice keypad using the index and middle finger of their dominant. The button assignment was kept constant across all participants.

The audiovisual stimuli were presented in two different conditions: matching visual and auditory input (*audiovisually congruent*) and uninformative visual input (*visually unspecific*). Each audiovisual condition was presented blockwise (similar to e.g. Klucharev et al., 2003; Winneke & Phillips, 2011), with the gaze directed to the left or right monitor, respectively, resulting in two blocks per audiovisual condition. Prior to each block, participants were informed about the condition and gaze direction. During a trial, two stimuli were simultaneously displayed on the left and right monitor, one of which always being a target (/yes/ or /no/) and the other one being a distractor (German number words from /one/ to /ten/). Stimuli in a trial were always uttered by two different talkers. In 80% of the trials, the location of the presented target matched the fixated monitor, in 20% of the trials it appeared on the other monitor. However, the present analysis focused on trials with fixated targets. One block consisted of 120 trials (i.e., 96 trials with targets at the fixated position and 24 trials with targets at the non-fixated position), resulting in a total of 480 trials (i.e., 384 trials with targets at the fixated position). The order of block presentation was counterbalanced to avoid sequence effects. Note, that the experiment was embedded in a study containing two sub-experiments (see Begau et al., 2021); both sub-experiments were counterbalanced in order. Between each block, participants had the possibility of a self-paced rest. No feedback was given during the experiment.

### 2.5 Lipreading assessment

We concluded the experimental session with a lipreading assessment, in which we presented 30 sentences (structure: name, verb, number, adjective, noun) spoken by a muted female talker. Details of the assessment are described in a previous study (Begau et al., 2021). Two tasks were given: A cue and a recognition task, with a maximum sum score of 30 points each. Prior to sentence presentation, two words were presented, one of which would appear in the sentence (cue task). After each sentence, participants were presented another word and had to indicate if it appeared in the sentence (recognition task). In the cue task, the mean sum score was 22.89 (*range* = 15 – 29, *SE* = 1.04) in the older and 24.05 (*range* = 12 – 30, *SE* = 1.06) in the younger group. In the recognition task, the mean sum score was 16.61 (*range* = 12 – 22, *SE* = 0.61) for the older and 17.67 (*range* = 13 – 27, *SE*= 0.75) for the younger group. Both groups did not differ in the recognition, *V* = 152.5, *p* = .295, *r* = .17 and the cue task, *t*(37) = -1.07, *p* = .293, *g* = -0.34, respectively.

### 2.6 EEG recording and preprocessing

#### 2.6.1 Recording

The continuous EEG signal was recorded from 64 Ag/AgCl electrodes (BrainCap; Brainvision, Gilching, Germany) with a sampling rate of 1000Hz (QuickAmp DC Amplifier, Brainvision, Gilching, Germany). Electrodes were evenly distributed across the scalp according to the extended international 10-20 system was used. The electrodes at AFz and FCz were used as online ground and reference, respectively. During electrode preparation, impedance was kept below an average of 10 kΩ.

#### 2.6.2 Preprocessing

For subsequent data preprocessing and analysis, MATLAB (2019a) with the toolboxes EEGLAB (version 14-1-2b, Delorme & Makeig, 2004) and ERPLAB (version v7.0 Lopez-Calderon & Luck, 2014) were used. A 0.5 Hz high-pass (6601 point, 0.5Hz transition bandwidth, 0.5Hz passband edge, 0.25Hz cutoff frequency) and 30 Hz low-pass Hamming windowed sinc FIR filter (441 point, 7.5Hz transition bandwidth, 30Hz passband edge, 33.75Hz cutoff frequency, using pop_eegfiltnew()) were applied to the continuous data. Subsequently, flat channels as well as channels contaminated by artifacts were removed. To identify the latter, kurtosis (using pop_rejchan(), normalized measure with SD = 5 as absolute threshold) and probability (same parameters as before) parameters were considered. Overall, an average of 4.66 channels were rejected (*range* = 1-9, *SE* = 0.33). Rejected channels were then spherically interpolated prior to re-referencing the signal to the average of all electrodes. Afterwards, the signal was epoched and stimulus-locked into sections from -2300 to 2400 ms relative to sound onset. The baseline period was set from -1800 to -1500 ms relative to sound onset. For further artifact rejection, we applied independent component analysis (ICA) to a subset of the data, high-pass filtered on 1 Hz (Hamming windowed sinc FIR, 3301 point, 1Hz transition bandwidth, 1Hz passband edge, 0.5Hz cut-off frequency), using every second trial and a sampling rate of 250 Hz for faster computation. Applying the algorithm ICLabel (Pion-Tonachini et al., 2019), all IC components received a probability estimate for the categories brain, muscle, eye, heart activity, line noise and other. IC weights and ICLabel results were transferred back to the whole dataset. ICs that received a probability estimate of less than 30% for the category brain or above 30% for the categories muscle, eye and heart activity, line noise or other activity were excluded (*M* = 34.97, *SE* = 1.31, *range* = 15 – 51). Finally, to remove remaining artefactual epochs, we applied an automatic trial rejection procedure (using the EEGLAB function pop_autorej()), resulting in the rejection of 86.46 trials on average (*SE* = 3.60, *range* = 33 – 134). The preprocessed data can be accessed via https://osf.io/.

### 2.7 Data Analysis

#### 2.7.1 Behavioral Analysis

We analyzed response times and accuracy data. Response times were measured relative to the end of fade-in in the target stimulus video, (i.e., at 500 ms post video onset). We chose this time point because it is a uniform starting point: otherwise, depending on the audiovisual condition and stimulus, responses could be influenced by lip movement and the latency between lip movement and sound onset. Only correct trials where the gaze direction matched the target location were analyzed. Trials with response times below or above three standard deviations from the mean were considered outliers and excluded from analysis (i.e., 5 trials in total).

#### 2.7.2 Time-Frequency analysis

For time-frequency analysis, we computed ERSPs separately for each age group and audiovisual condition. We chose only trials where the gaze direction matched the target location. Additionally, only trials with correct responses were included. This results in an average of 83.03 trials (*SE* = 0.92) for the audiovisually congruent and 81.21 trials (*SE* = 0.81) for the incongruent condition. For the analysis of event-related theta-band perturbations, frequencies between 4 and 8 Hz (Peelle & Sommers, 2015) at a frontal electrode cluster (Fz, FCz, Cz, FC1 and FC2) were considered. Please note, while Morís Fernández et al., 2018, decided for a cluster around Cz we decided for a more frontal cluster, taking into account aging-related frontal shifts (see for example Davis, Dennis, Daselaar, Fleck, & Cabeza, 2008). Event-related alpha-band perturbations (8 to 12 Hz) were analyzed at the same frontocentral cluster as well as a parietooccipital cluster (POz, Pz, P1 and P2), taking previous work by Keller and colleagues (2017) and Misselhorn and colleagues (Misselhorn et al., 2019)as orientation. Finally, event-related beta-band perturbations(16 to 30 Hz) were investigated at a central electrode cluster (Cz, CPz, FCz, C1, C2), since beta power modulations in audiovisual stimuli have been demonstrated in frontal, central and parietal locations (e.g. Roa Romero et al., 2015; Schepers et al., 2013). We performed Morlet wavelet convolution on epoched data over 52 frequencies from 4 to 30 Hz, starting with 3 cycles at the lowest frequency, with steps of 0.5 cycles, ending at 11.25 Hz for the highest frequency, generating 200 time points for each ERSP. A time window of 300ms for a dB spectral baseline was chosen between -400 and -100 ms prior to sound onset. This period corresponds with the ISI, that is participants were presented with black screens during that time. A condition specific baseline was used. Prior to baseline removal a single trial normalization over the full trial length was conducted (see Grandchamp & Delorme, 2011).

#### 2.7.3 Statistical Analyses

For the analysis of response times, we conducted a 2 (age; old vs. young) by 2 (audiovisual condition; audiovisually congruent vs. visually unspecific) mixed ANOVA. Normal distribution was inspected using Shapiro-Wilk test (Shapiro & Wilk, 1965). In case of violation, we calculated nonparametric ANOVA-like statistics using the R package nparLD (Noguchi et al., 2012). Note, that here, the denominator degree of freedom is set to infinite for within subject effects and thus not reported. If possible, effect sizes were calculated, that is, *η*_*p*_^*2*^ for ANOVA and Hedge’s *g* for t-tests (Lakens, 2013). For accuracy data, we conducted a 2 (age; old vs. young) by 2 (audiovisual condition; visually unspecific vs, audiovisually congruent) mixed ANOVA. Since the data showed clear ceiling effects, non-parametric analyses were chosen. For follow-up comparisons, Wilcoxon *U* test for dependent samples was applied for a comparison of audiovisual conditions and Wilcoxon’s *U* for independent samples for age group comparisons (Wilcoxon, 1946). For Wilcoxon, *r* was reported as an effect size estimate (Lakens, 2013).

#### 2.7.2 Time-Frequency Data

We compared event-related spectral perturbations in the theta, alpha and beta band across audiovisual conditions and age groups. Note, that the term “power” in that case refers to spectral power changes relative to a pre-stimulus baseline (Makeig et al., 2004). In accordance with previous literature (Morís Fernández et al., 2018), we expect audiovisual condition differences in the mean peak of theta enhancement immediately after sound onset, as revealed by the respective ERSPs. Therefore, we computed a 2 (old vs. young) by 2 (visually unspecific vs. audiovisually congruent) mixed ANOVA, using the mean peak amplitude as dependent variable. The mean peak was calculated 100ms around the local maximum within a time window between 0 and 700 ms. Cluster permutation analysis (Maris & Oostenveld, 2007) was used for statistical testing of the alpha and beta band, since we did not specifically expect peak differences. That is, for beta power previous literature indicated audiovisual condition differences to arise before and after the peak of power suppression (e.g. Roa Romero et al., 2015). For this, ERSPs were averaged over selected frequency bands and electrode clusters. For ERSPs time-locked to sound onset, we chose an analysis time window of interest from 0 to 700 ms, that is from the auditory onset to the end of the lip movement. We calculated t-tests for each time-point in the original data. Specifically, independent samples t-tests were used for the main effect age (e.g., young vs. old) and the interaction (e.g., difference of congruent vs. unspecific in the old vs. in the young group) and dependent samples t-test for the main effect audiovisual information (e.g., congruent vs. unspecific). Clusters were formed with adjacent voxels reaching a significance threshold of *p* <.05 or *p* <.01, respectively, were used to estimate the maximal cluster size under the null hypothesis, condition labels were shuffled (i.e., ERSPs were randomly assigned to a given audiovisual task condition) across 50’000 permutations and respective t-tests were computed for each time point in the permuted data. For this, voxels within the permuted data had to reach a significance threshold of *p* < .05. For original data clusters to reach significance, they had to surpass the 95th and 99th percentile of the permuted maximum cluster size, respectively.

## 3 Results

### 3.1 Behavioral Analysis

#### 3.1.1 Response times

There were longer response times in the older compared to the younger group, *F*(1,36.31) = 4.17, *p* = .049. Moreover, congruent stimuli elicited shorter response times than incongruent stimuli, *F*(1, ∞) = 55.43, *p* < .001. An interaction of both factors was not present, *F*(1, ∞) = 0.75, *p* = .385 (see fig. 2).

**Figure 2:**
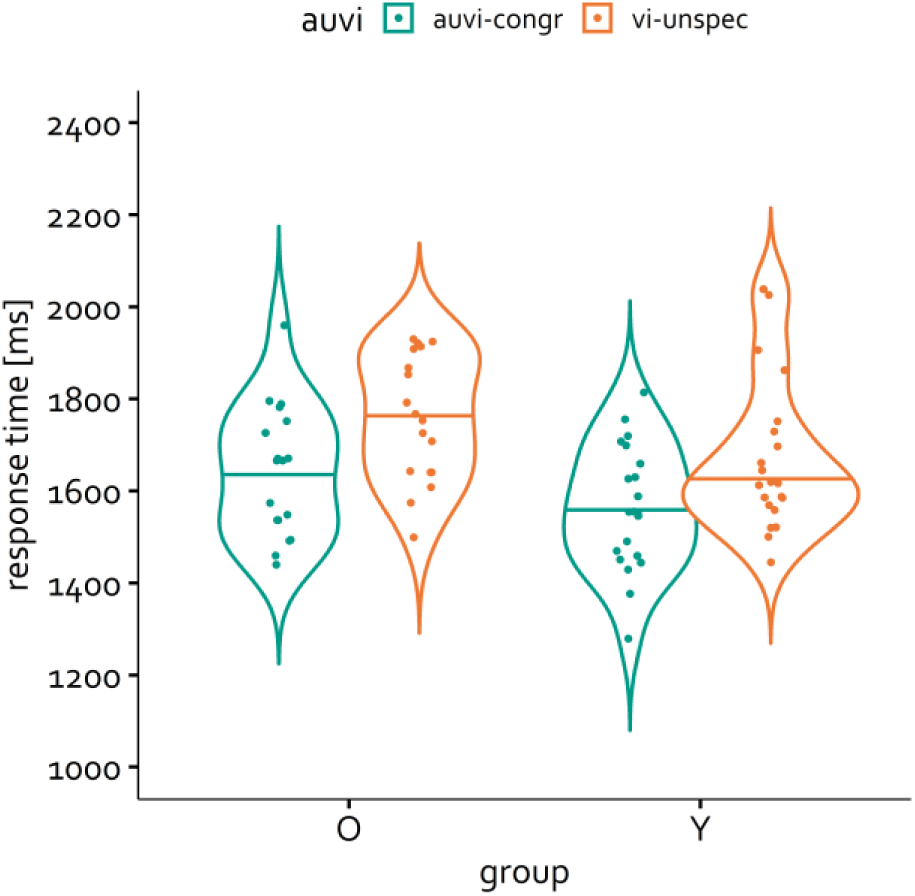
Violin plots of response times (mean and individual values) for audiovisual congruent stimuli (auvi-congr) and audiovisual stimuli containing only uninformative visual information (vi-unspec), and for younger (Y) and older (O) participants. Response times (ms) were calculated relative to the end of video fade-in (i.e., 500ms post video-onset and 1000ms prior to sound onset) for correct trials only. Main effect age p <.05, main effect audiovisual condition p <.001.

#### 3.1.2 Accuracy

Overall, participants showed good performance on the task, which was visible in accuracy measures reaching ceiling level (fig. 3). Statistical analysis showed no difference between the age groups, *F*(1,36.97) = 0.08, *p* = .776, but higher accuracy for stimuli with congruent compared to incongruent information, *F*(1, ∞) = 8.58, *p* = .003. No interaction was found, *F*(1, ∞) = 1.10, *p* = .189.

**Figure 3:**
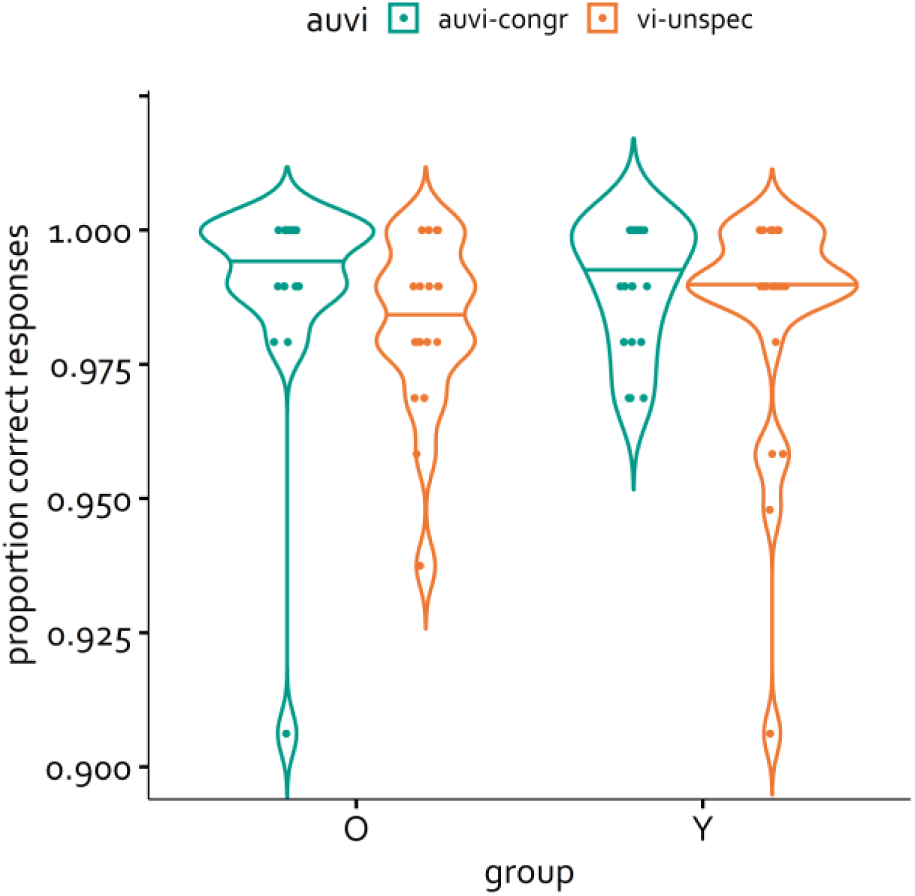
Violin plots for proportion of correct responses (mean and individual values) for audiovisual congruent stimuli (auvi-congr) and audiovisual stimuli containing only uninformative visual information (vi-unspec), and for younger (Y) and older (O) participants. Main effect audiovisual condition *p* <.01.

### 3.2. ERSP Analysis

#### 3.2.1. Theta oscillations

Figure 4 shows the development of theta perturbations (4-8 Hz) after sound onset at a frontocentral electrode cluster including FCz, Cz, Fz, FC1, and FC2. A first peak in theta perturbations right after sound onset can be observed in all conditions. Analyzing the mean theta peak, we found no general differences between both groups, *M*_*young*_ =1.33 dB, *SE*_*young*_ = 0.12, *M*_*old*_ = 1.59 dB, *SE*_*old*_ = 0.17, *F*(1, 37) = 1.19, *p* = .282, *η*_*p*_^*2*^ < .03. Amplitudes for visually unspecific information were larger than for congruent stimuli, *M*_*Vunspec*_ = 1.75 dB, *SE*_*Vunspec*_ = 0.15, *M*_*AVcongr*_ = 1.15 dB, *SE*_*AVcongr*_ = 0.12, *F*(1, 37) = 25.20, *p* < .001, *η*_*p*_^*2*^ = .41. We found an interaction of both factors, *F*(1, 37) = 6.99, *p* = .012, *η*_*p*_^*2*^ = .16, showing that the difference between audiovisual conditions was driven by the older group, *t*(17) = 6.22, *p* <.001, *g* = 1.40, while there was no difference within the younger group, *t*(20) = 1.71, *p* = .103, *g* = 0.36.

**Figure 4:**
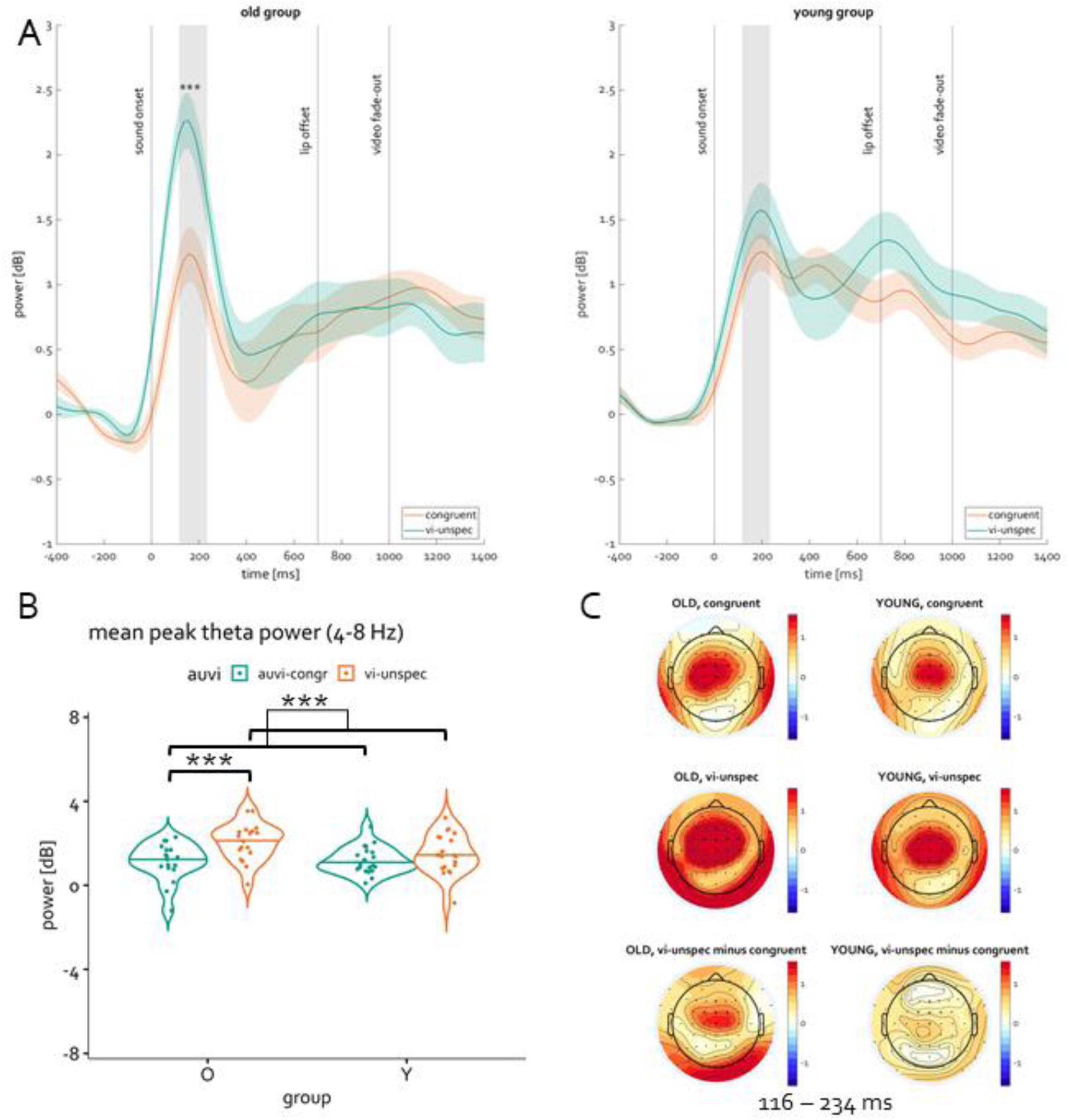
Mean event-related theta perturbations (4 – 8 Hz) over frontocentral electrode cluster, shown separately for audiovisual congruent stimuli (auvi-congr) and audiovisual stimuli containing only uninformative visual information (vi-unspec), and for younger (Y) and older (O) participants. (A) Event-related theta perturbations relative to the onset of acoustic speech. The grey area indicates mean peak area (116 – 234 ms), the shaded areas around the line plots indicate the standard error of the mean. (B) Violin plots (mean and individual values; ***: p < .001). (C) Topographies of theta perturbations for audiovisual congruent and audiovisual stimuli containing only uninformative information (vi-unspec) as well as visually unspecific-minus-congruent difference topographies averaged across the time window marked in (A). Clustered electrodes were FCz, Fz, Cz, FC1, FC2.

#### 3.2.2 Alpha Oscillations

In the development of event-related alpha perturbations from sound onset to lip movement offset (fig. 5), we observed an initial increase in frontal to central electrodes, followed by a suppression in alpha power. This suppression could also be observed at occipito-parietal sites (fig. 6). As can be seen in figure 6, alpha power suppression as indicated by ERSPs peaked around 400 ms, once again followed by an increase in alpha power. Condition and group differences were then compared in both a fronto-central and a parieto-occipital electrode cluster, the significance threshold was Bonferroni corrected to *p*= .025 and *p* = .005, respectively (as explained in e.g. Luck, 2014). In the parieto-occipital cluster (Pz, POz, P1, P2), cluster-based permutation differences revealed no significant difference between age groups, audiovisual conditions, or the double difference between audiovisual conditions across both age groups (all *p* > .05). The frontocentral cluster did not reveal any differences between age groups and audiovisual conditions either (all *p* >.05).

**Figure 5:**
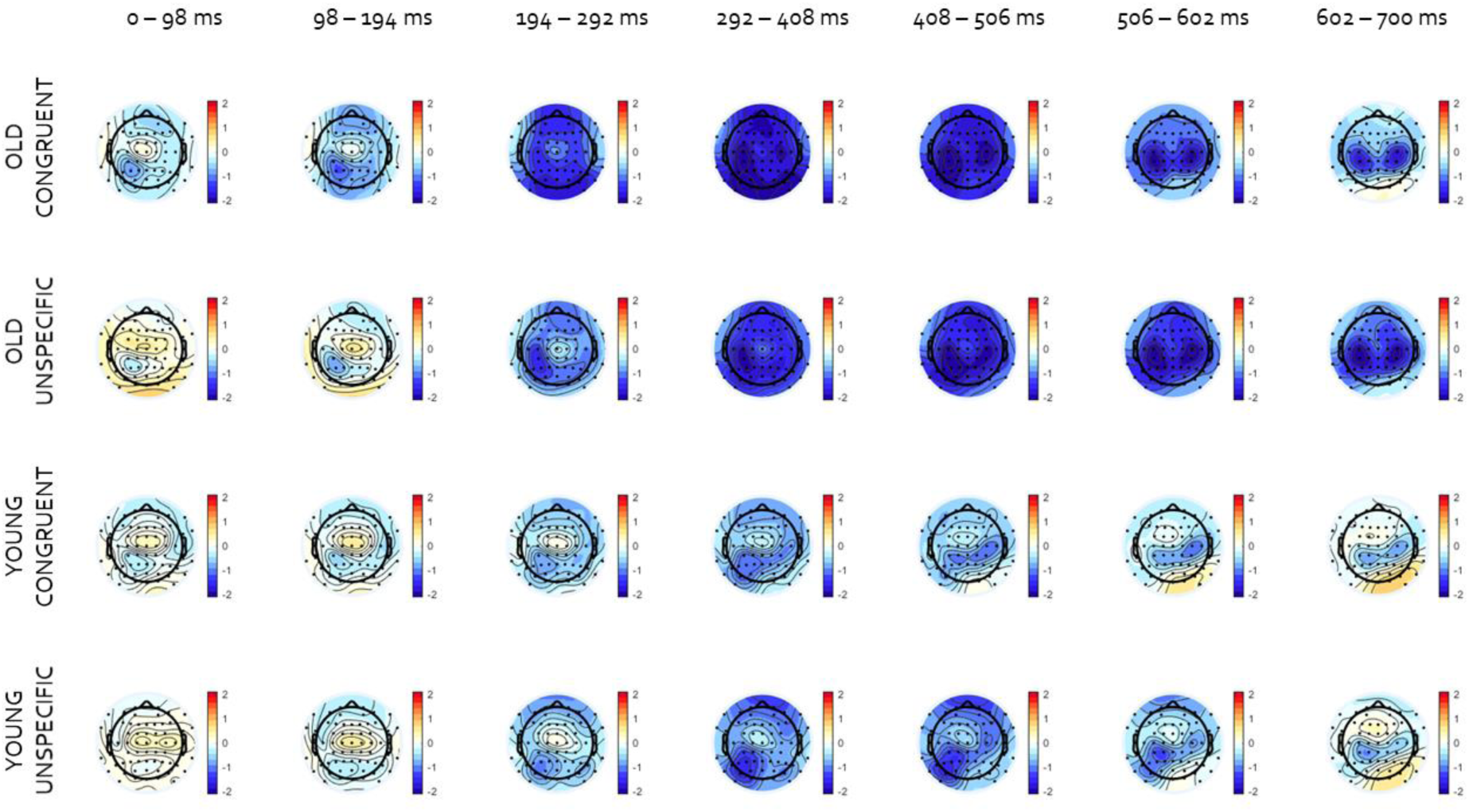
Event-related alpha perturbations (ERSPs) (8 – 12 Hz) topographies from sound onset to lip movement offset for audiovisual congruent stimuli and visually unspecific stimuli, and for younger and older participants: ERSPs were averaged over indicated time-windows. Sound onset was at 0 ms, average lip movement offset around 700 ms.

**Figure 6:**
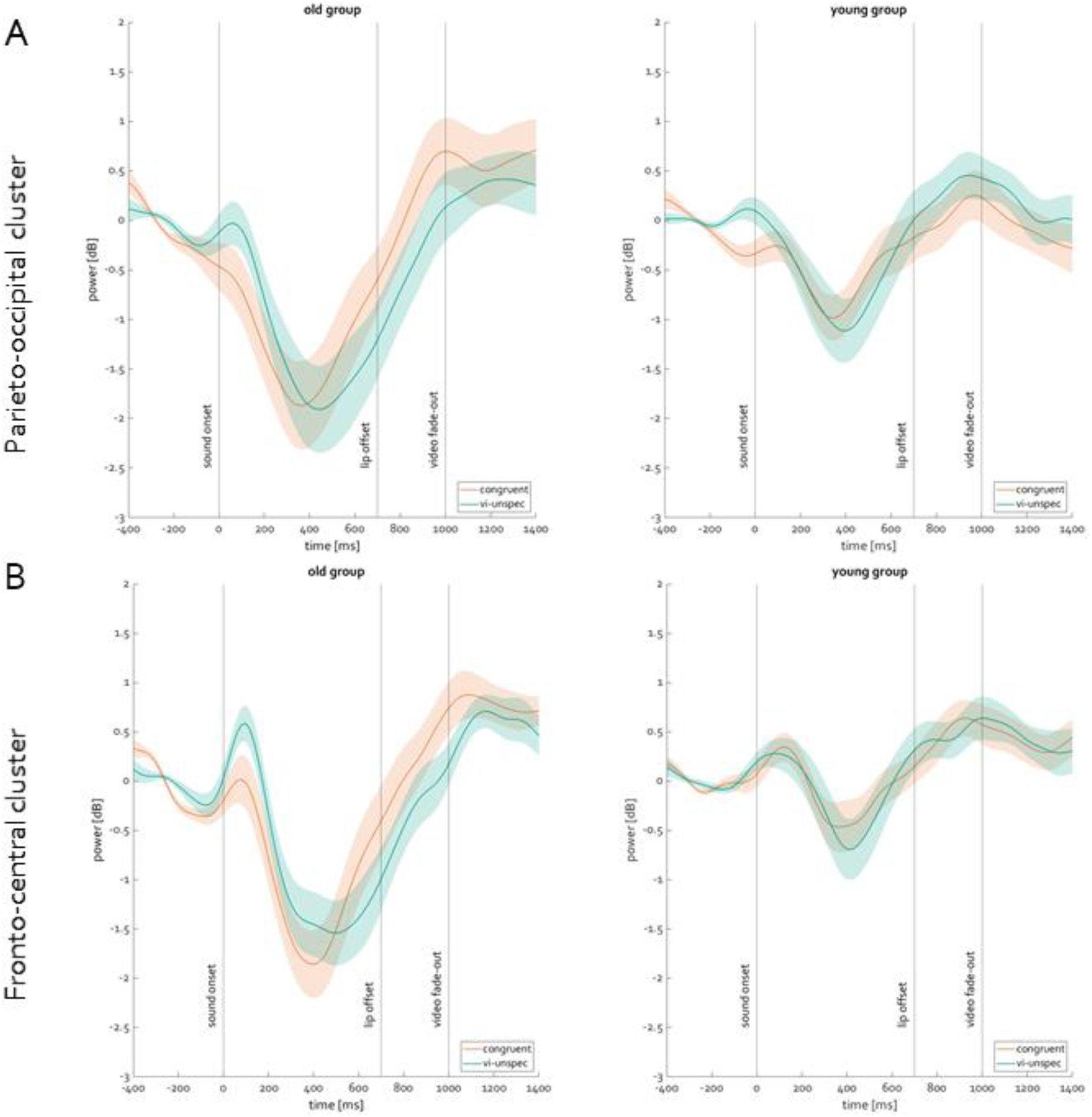
Mean event-related alpha perturbations (8 – 12 Hz) over parieto-occipital (A) and fronto-central (B) electrode cluster relative to the onset of acoustic speech for audiovisual congruent stimuli (congruent) and audiovisual stimuli containing only uninformative visual information (vi-unspec), and for younger and older participants: Electrode cluster (A) Pz, POz, P1, P2, (B) FCz, Cz, Fz, FC1, and FC2. The shaded areas around the line plots indicate the standard error of the mean.

#### 3.2.3 Beta oscillations

Beta oscillations were analyzed at a central electrode cluster (Cz, CPz, FCz, C1, C2). Figure 7A shows event-related beta perturbations, revealing a general decrease in beta power after sound onset, that peaks around 390 ms. Subsequently, beta power increases again, reaching overall higher levels by the end of stimulation. Notably, around sound onset there is a slight increase in beta power for visually unspecific stimuli before a strong suppression can be observed. This initial positive peak does not occur with audiovisually congruent stimuli. Cluster permutation analyses revealed a significant cluster (156-700ms) for the age group difference (p <.01), which is around the time of maximum beta suppression. As depicted in figure 7A, ERSPs reveal a beta power suppression that is stronger in older compared to younger participants. There was no difference between audiovisual conditions across both groups (p > .05), however we found a significant cluster (0 – 194 ms) for an interaction of audiovisual condition and age group (p <.05). This cluster occurs during sound onset and the following suppression in beta power, as revealed by ERSPs. Subsequently, we ran cluster permutation tests for both groups separately and found a difference between the audiovisual conditions only in the older (cluster from 0 to 214 ms, p <.05), but not in the younger group (p > .05). Figure 7B shows that event-related beta perturbations reveal a stronger initial suppression of beta power in audiovisually congruent compared to visually unspecific stimuli in the older group.

**Figure 7:**
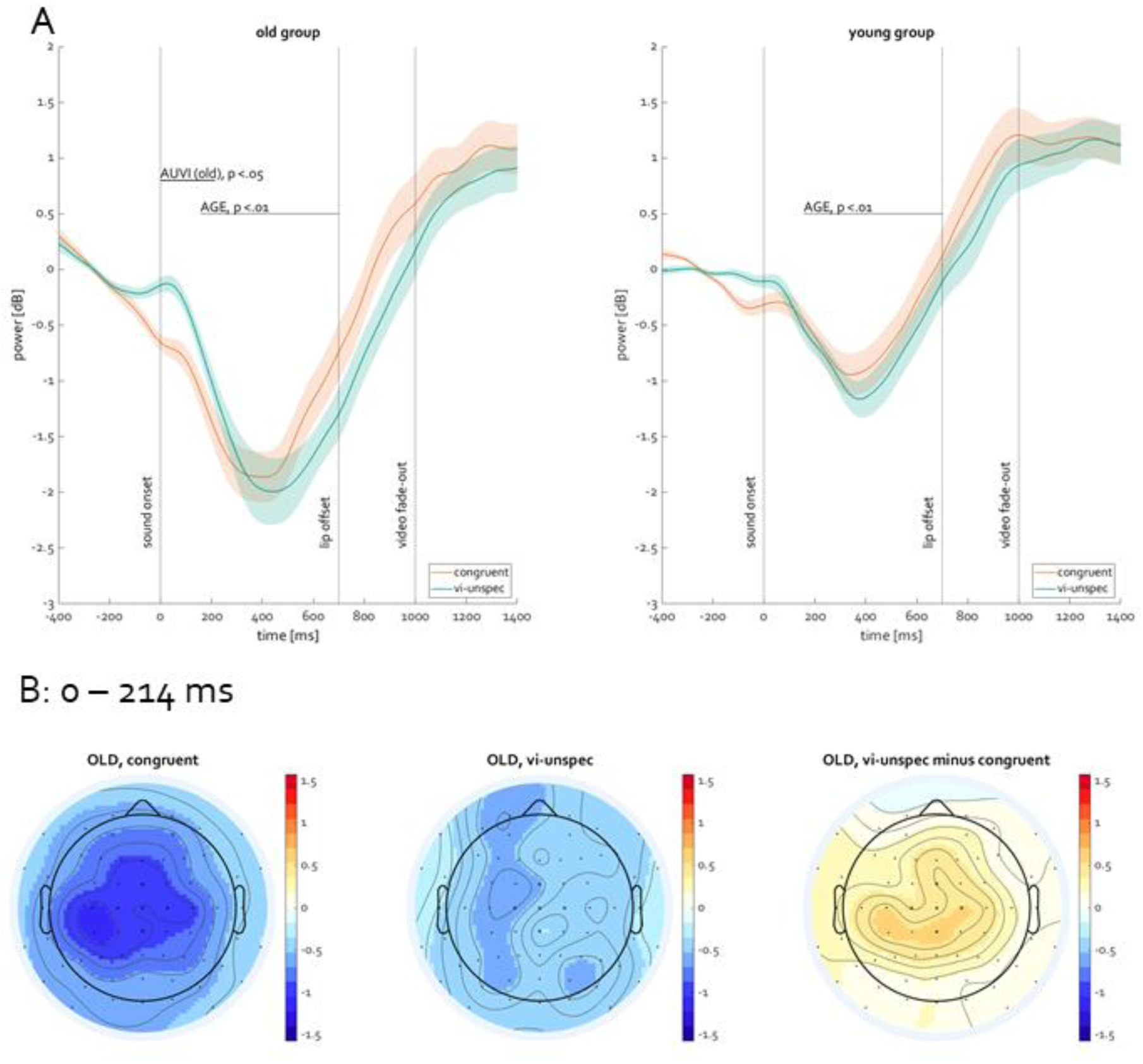
Mean event-related beta perturbations (16 – 30 Hz) over central electrode cluster: (A) Mean event-related beta perturbations relative to the onset of acoustic speech for audiovisual congruent stimuli (congruent) and audiovisual stimuli containing only uninformative visual information (vi-unspec), and for younger and older participants. Clustered electrodes were Cz, CPz, FCz, C1, C2. The shaded areas around the line plots indicate the standard error of the mean. (B) Topographies of event-related beta perturbations of the older group for audiovisual congruent and audiovisual stimuli containing only uninformative visual information (vi-unspec) as well as visually unspecific-minus-congruent difference topographies averaged over the time window of audiovisual condition difference in older group (0 to 214 ms): Cluster permutation analysis was conducted for the time between sound onset (0ms) and the end of lip movement (700 ms). Significant clusters for pairwise comparisons are indicated by lines.

## 4 Discussion

In this study, we investigated the processing of audiovisual speech stimuli in younger and older adults, where the visual speech information either matched the auditory input (i.e., audiovisually congruent) or did not provide any task-relevant information (i.e., visually unspecific). We hypothesized that the processing of audiovisually congruent speech would be facilitated compared to that of visually unspecific speech. This was expected to be visible in performance measures, such as response time and accuracy, as well as theta, alpha and beta oscillations. To sum up, responses were faster and more accurate for audiovisually congruent compared to incongruent stimuli. Behaviorally, age differences were only found in response times, with slower responses in the older group. In event-related spectral perturbations (ERSPs), we found a larger mean peak amplitude in frontocentral theta power for visually unspecific compared to audiovisually congruent stimuli. Importantly, this difference was only present in the older group. Moreover, no modulations in event-related alpha perturbations across conditions were found, neither at parietooccipital, nor at frontocentral electrode locations. For event-related beta perturbations, we observed a generally stronger suppression at central electrodes in the older compared to the younger group. Only in the older group, we found an early cluster of stronger beta suppression for congruent compared to visually unspecific stimuli. These results are discussed in detail in the following.

### 4.1 Processing of audiovisual information

Behaviorally, participants showed very good task performance, which led to ceiling effects in response accuracy. While this could be expected due to the relatively easy task, faster response times and higher accuracy were found across both age groups for audiovisually congruent information. This behavioral audiovisual benefit is in line with previous studies showing better performance for congruent audiovisual compared to unisensory information (Sommers et al., 2005; van Wassenhove et al., 2005; Winneke & Phillips, 2011). Our finding also replicates findings from a previous study adopting analogous modulations of audiovisual informational content (Begau et al., 2021). In correspondence to these behavioral results, we found differences in the theta and beta band perturbations in the EEG between audiovisual conditions as well as between younger and older participants. Note again, that the discussed power differences in our results refer to changes in spectral perturbations relative to a pre-stimulus baseline.

For event-related theta perturbations, a stronger increase in theta power for unspecific visual information compared to congruent information. This difference was only present in the older group. In audiovisual stimuli, an increase in theta power has been linked to incongruence processing and the need for cognitive control to successfully enable audiovisual integration (Michail et al., 2021 using the sound induced flash illusion; Morís Fernández et al., 2015 using McGurk stimuli). At the time of sound onset, the preceding visual information is already partially processed. Thus, the formation of a prediction of the cooccurring auditory information is enabled, leading to a facilitated processing of the auditory information. In case of visually unspecific information, no prediction can be made, thus the processing of the auditory stimulus cannot be facilitated by preceding visual stimulation. This is in line with the analysis-by-synthesis (van Wassenhove et al., 2005) or predictive coding hypotheses (Arnal & Giraud, 2012). Furthermore, even though the unspecific visual information does not contain task-relevant speech information, the comparison of visual and auditory information would still lead to the detection of an audiovisual mismatch. Accordingly, an increase (fronto-)central theta has previously been interpreted as the activation of a conflict processing network (Morís Fernández et al., 2018) following the detection of a mismatch between expected and actual auditory information in audiovisual speech stimuli. Overall, our finding is in line with other studies demonstrating frontocentral theta increase to be stronger in audiovisual compared to unimodal stimuli (Keller et al., 2017), McGurk illusions (Keil et al., 2012; Morís Fernández et al., 2018), as well as temporal audiovisual asynchrony (Simon & Wallace, 2018).

Meanwhile, looking at event-related beta perturbation, we found a suppression in central beta power shortly after the theta increase, with a stronger suppression in congruent compared to visually unspecific stimuli following sound onset. This stronger beta suppression was found also only in the older group. This suppression in beta power (as shown by ERSPs) likely reflects the processing of the auditory stimulus, in accordance with the notion that beta power suppression is linked to language processing (Weiss & Mueller, 2012). Moreover, decreased activity in the beta band has been associated with stimulus-driven rather than top-down processing (Engel & Fries, 2010). This may be especially important in audiovisual speech stimuli, where the auditory usually follows the visual information. Accordingly, since the auditory information is task-relevant in the present study and resolves any ambiguity arising from the visual speech, the comparison of the predicted and actual auditory information is required, as suggested by previous studies on predictive coding (Arnal & Giraud, 2012). Furthermore, beta power suppression has been shown to reflect integration processes in audiovisual stimuli (Keil & Senkowski, 2018). Previous findings further link early beta suppression to the fusion of audiovisual stimuli, while later beta suppression is associated with top-down integration (Michail et al., 2021; Roa Romero et al., 2015).

Taken together, while a benefit of congruent audiovisual information can be observed behaviorally, both event-related theta and beta perturbations reflect the neural processing of the audiovisual stimuli. While theta is likely sensitive to the early detection of audiovisual mismatch and the initiation of further processes that require cognitive control, beta seems to be linked to processes linked to the integration of the multisensory percept, monitoring the sensory input. This is in line with the notion that different cortical oscillations are associated with different stages of audiovisual speech processing, as suggested by the framework of Keil and Senkowski (2018). There, the authors argue theta to be involved in feedback feed forward loops in case of audiovisual incongruence, while beta is thought to be involved in the comparison of the auditory and visual information.

### 4.2 The influence of the multi-talker scenario

While audiovisual speech integration and processing is often investigated using only one talker (e.g. Fingelkurts et al., 2007; Morís Fernández et al., 2018; Roa Romero et al., 2015), we presented a simplified multi-talker scenario, in which the listener had to suppress the concurrent auditory information of the second talker. The engagement of additional attentional resources to focus on the target talker has therefore been expected, as indicated by ERSPs showing a suppression of alpha power after sound onset over parieto-occipital and fronto-central electrode sites. In multisensory attention paradigms, posterior alpha oscillations have been linked to selective attention mechanisms complementary to theta as a marker for divided attention (Keller et al., 2017). Furthermore, frontal alpha suppression has been linked to top-down regulation of perceptual gain, while parieto-occipital alpha was interpreted as a marker for intersensory orienting (Misselhorn et al., 2019). Contrary to our expectations, the magnitude of alpha suppression (measured as ERSPs) did not differ between conditions and groups. This could be due to the fact that both conditions required selective attention to a similar extent. In both the audiovisually congruent and visually unspecific stimuli, auditory information provided the most reliable task-relevant information. Due to the multi-talker setting, the auditory information was also susceptible to interference by the simultaneously presented distractor, while the competing visual information was visible only peripherally. Thus, participants may have similarly directed their (intersensory) attention towards the auditory input and selectively enhanced the perceptual gain from the targeted auditory information in both audiovisual conditions.

Previous studies demonstrated the association of parieto-occipital alpha power with the suppression of visual speech in incongruent audiovisual speech stimuli (O’Sullivan et al., 2019), or unreliable (i.e., blurred) visual speech stimuli (Shatzer et al., 2018). However, Shatzer and colleagues (2018) did not present a multi-talker setting and found the strongest differences in alpha suppression in more lateral parietooccipital sites rather than around the midline. While O’Sullivan and colleagues (2019) presented a multi-talker paradigm, the distraction arising from the visual stimulus may be stronger than in the present study. There, participants needed to either actively attend or ignore the auditory input that belonged to the fixated visual information, which probably resulted in a higher salience of the visual information and thus in much more distracting concurrent visual information when the incongruent auditory input was task relevant. In contrast, visually unspecific information in the present study might not be actively distracting since it does not provide any misleading information.

The presence of more than one talker may also have an influence on audiovisual speech integration reflected by event-related beta band perturbations. In the present study, an early beta suppression difference was found between conditions. However, this effect appeared only in the older group, as will be discussed in section 4.3. Inversely to previous single-speaker studies (e.g., Roa Romero et al., 2015), this suppression was stronger in the congruent condition. Presenting the stimuli in a multi-talker setting, probably made audiovisual integration more demanding. In accordance, beta suppression has shown to be stronger with higher memory load (Michail et al., 2021) or enhanced noise levels (Schepers et al., 2013) during stimulus presentation. Another study demonstrated stronger beta suppression for mismatched versus matched gesture-speech combinations in clear speech, but not in degraded speech (Drijvers et al., 2018). In the latter study, higher beta power was found for mismatched compared to matched combinations. Therefore, the quality of the speech input may have an impact on beta power modulation, visible in stronger beta suppression in the presence of unambiguous, clearly distinguishable information. Overall, the cognitive demands due to the presence of multiple valid information streams in the multi-talker setting with audiovisually congruent stimuli may have been higher than the audiovisual mismatch arising from visually unspecific information. Since the conditions were presented block wise, less resources for integration may have been activated. While studies investigating McGurk illusions (Michail et al., 2021; Roa Romero et al., 2015) argue that early beta suppression reflects audiovisual fusion in illusory stimuli, it may also reflect a general binding mechanism, superficially associating the talker’s face to the according auditory information.

### 4.3 Processing differences due to aging

The present study also investigated age-related differences in processing audiovisual speech in a multi-talker scenario. General age differences were only found in larger response times in the older group, but not in accuracy measures, as well as in stronger beta suppression, as shown by ERSPs, in the older group. The behavioral differences are well in line with a general speed-accuracy trade-off in older adults (Salthouse, 1979). Thus, high accuracy in the older adults comes at the cost of larger response times. The slower responses can also be explained by the generalized slowing in older adults leading to a slower processing of sensory information (Salthouse, 2000). The relatively low task difficulty may explain, why unlike previous studies (Begau et al., 2021; Sekiyama et al., 2014; Winneke & Phillips, 2011), we did not find a greater audiovisual benefit for older adults.

We did not find general age differences in event-related theta perturbations, which contrasts earlier findings, such as a study by Cummins and Finnigan (2007) demonstrating lower theta power in older adults. While the authors presented a retention and recognition task, our participants only had to do a relatively easy detection task, possibly explaining the differing results. This is further supported by findings of increased midfrontal theta activity with higher task difficulty and higher cognitive load (Cavanagh & Frank, 2014; Maurer et al., 2015). Since the task itself (detection of the target word) was relatively easy, as can be seen in the overall high accuracy ceiling effects, we may not find general age differences. Another aspect that could have an impact here is that the importance of perceptual input changes with age (Habak et al., 2019). Specifically, previous research demonstrates that older adults rely more strongly on visual speech input in audiovisual scenarios (Sekiyama et al., 2014). In task blocks with uninformative visual information this might turn out to be a drawback; while younger participants may have opted to ignore the uninformative visual input for task completion (due to the blocked design of the study), older participants may have failed to do so. This may also explain, why contrary to our expectations and previous findings (Enriquez-Geppert & Barceló, 2018), event-related beta perturbations revealed a stronger suppression in the older than in the younger group. Hence, one could speculate that this stronger suppression may be linked to more activated resources to integrate audiovisual information in an attempt to entirely process the given information, whether it is task-relevant or not. This may also explain, why differences between audiovisual conditions in the theta and beta band were only observable in the older group, while there were no differences in the younger group. In line with the inhibition deficit hypothesis (e.g. Rey-Mermet & Gade, 2018; Stothart & Kazanina, 2016), older adults may need to resolve the conflict arising from the audiovisual mismatch, while younger adults learn to successfully ignore the visual information.

### 4.4 Conclusion

To summarize, the present study investigated audiovisual speech processing in a multi-talker scenario using realistic, ecologically valid stimuli. Thus, we provided further insights on the processing of natural speech in a complex listening environment. The present findings suggest that, while congruent audiovisual speech is beneficial for both older and younger adults on a behavioral level, we found age-related differences in the processing of audiovisual speech. We compared the processing and integration of audiovisual speech with congruent and uninformative visual speech and showed differences in theta enhancement and beta suppression after sound onset only in the older, but not in the younger group. This further emphasizes the importance of congruent visual speech for audiovisual speech information to be beneficial especially for older adults. For this age group, we provided further evidence for the association of increased frontocentral theta perturbations and an early conflict detection mechanism, as well as suppression in central event-related beta perturbations and more general integrative processes.

## Acknowledgements

The authors are grateful to Nina Abich, Christele Motcho, and Tobias Blanke for technical support and to Stefan Weber, Kristin Koberzin, Vera Birgel and Denise Böhle for running the experiment. They would also like to thank Kimberly Freytag, Bianca Zickerick and Nina Abich for volunteering as stimulus material.

## Conflict of interest statement

All authors disclose no actual or potential conflicts of interest including any financial, personal, or other relationships with other people or organizations that could inappropriately influence (bias) their work.

## Data availability statement

The data that support the findings of this study can be found via [OSF](https://osf.io/hm93t/?view_only=e86beac05d5b4ed08e42866b4fe65bb1) and will be made public after acceptance.

## Author contributions

The study was designed by Stephan Getzmann, Alexandra Begau, Daniel Schneider and Edmund Wascher. Alexandra Begau supervised data collection and conducted data analysis supported by support and helpful discussions by Laura-Isabelle Klatt. Results and interpretations were intensely discussed by all authors. Alexandra Begau drafted the manuscript, which was revised by all authors.

## Ethical statement

The study conformed to the Code of Ethics of the World Medical Association (Declaration of Helsinki) and was approved by the local Ethical Committee of the Leibniz Research Centre for Working Environment and Human Factors, Dortmund, Germany.

## Funding

This work was supported by a grant from the Deutsche Forschungsgemeinschaft (GE 1920/4-1).

## Notes

https://osf.io/hm93t/?view_only=e86beac05d5b4ed08e42866b4fe65bb1

### Competing Interest Statement

The authors have declared no competing interest.

## Literature

Arnal, L. H., & Giraud, A.-L. (2012). Cortical oscillations and sensory predictions. Trends in Cognitive Sciences, 16(7), 390–398. https://doi.org/10.1016/j.tics.2012.05.003

Baart, M., Stekelenburg, J. J., & Vroomen, J. (2014). Electrophysiological evidence for speech-specific audiovisual integration. Neuropsychologia, 53(1), 115–121. https://doi.org/10.1016/j.neuropsychologia.2013.11.011

Begau, A., Klatt, L.-I., Wascher, E., Schneider, D., & Getzmann, S. (2021). Do congruent lip movements facilitate speech processing in a dynamic audiovisual multi-talker scenario? An ERP study with older and younger adults. Behavioural Brain Research, 412, 113436. https://doi.org/10.1016/j.bbr.2021.113436

Bregman, A. S., & McAdams, S. (1994). Auditory Scene Analysis: The Perceptual Organization of Sound. The Journal of the Acoustical Society of America, 95(2), 1177–1178. https://doi.org/10.1121/1.408434

Bronkhorst, A. W. (2015). The cocktail-party problem revisited: early processing and selection of multi-talker speech. Attention, Perception, & Psychophysics, 77(5), 1465–1487. https://doi.org/10.3758/s13414-015-0882-9

Carson, N., Leach, L., & Murphy, K. J. (2018). A re-examination of Montreal Cognitive Assessment (MoCA) cutoff scores. International Journal of Geriatric Psychiatry, 33(2), 379–388. https://doi.org/10.1002/gps.4756

Cavanagh, J. F., & Frank, M. J. (2014). Frontal theta as a mechanism for cognitive control. Trends in Cognitive Sciences, 18(8), 414–421. https://doi.org/10.1016/j.tics.2014.04.012

Chandrasekaran, C., Trubanova, A., Stillittano, S., Caplier, A., & Ghazanfar, A. A. (2009). The Natural Statistics of Audiovisual Speech. PLoS Computational Biology, 5(7), e1000436. https://doi.org/10.1371/journal.pcbi.1000436

Cherry, E. C. (1953). Some Experiments on the Recognition of Speech, with One and with Two Ears. The Journal of the Acoustical Society of America, 25(5), 975–979. https://doi.org/10.1121/1.1907229

Cienkowski, K. M., & Carney, A. E. (2002). Auditory-Visual Speech Perception and Aging. Ear and Hearing, 23(5), 439–449. https://doi.org/10.1097/00003446-200210000-00006

Correa-Jaraba, K. S., Cid-Fernández, S., Lindín, M., & Díaz, F. (2016). Involuntary Capture and Voluntary Reorienting of Attention Decline in Middle-Aged and Old Participants. Frontiers in Human Neuroscience, 10(129), 1–13. https://doi.org/10.3389/fnhum.2016.00129

Cummins, T. D. R., & Finnigan, S. (2007). Theta power is reduced in healthy cognitive aging. International Journal of Psychophysiology, 66(1), 10–17. https://doi.org/10.1016/j.ijpsycho.2007.05.008

Davis, S. W., Dennis, N. A., Daselaar, S. M., Fleck, M. S., & Cabeza, R. (2008). Que PASA? The Posterior-Anterior Shift in Aging. Cerebral Cortex, 18(5), 1201–1209. https://doi.org/10.1093/cercor/bhm155

Delorme, A., & Makeig, S. (2004). EEGLAB: an open source toolbox for analysis of single-trial EEG dynamics including independent component analysis. Journal of Neuroscience Methods, 134(1), 9–21. https://doi.org/10.1016/j.jneumeth.2003.10.009

Drijvers, L., Özyürek, A., & Jensen, O. (2018). Alpha and Beta Oscillations Index Semantic Congruency between Speech and Gestures in Clear and Degraded Speech. Journal of Cognitive Neuroscience, 30(8), 1086–1097. https://doi.org/10.1162/jocn_a_01301

Engel, A. K., & Fries, P. (2010). Beta-band oscillations—signalling the status quo? Current Opinion in Neurobiology, 20(2), 156–165. https://doi.org/10.1016/j.conb.2010.02.015

Enriquez-Geppert, S., & Barceló, F. (2018). Multisubject Decomposition of Event-related Positivities in Cognitive Control: Tackling Age-related Changes in Reactive Control. Brain Topography, 31(1), 17–34. https://doi.org/10.1007/s10548-016-0512-4

Fingelkurts, A. A., Fingelkurts, A. A., & Krause, C. M. (2007). Composition of brain oscillations and their functions in the maintenance of auditory, visual and audio–visual speech percepts: an exploratory study. Cognitive Processing, 8(3), 183–199. https://doi.org/10.1007/s10339-007-0175-x

Friedman, D. (2011). The Components of Aging. In E. S. Kappenman & S. J. Luck (Eds.), The Oxford Handbook of Event-Related Potential Components (Vol. 1). Oxford University Press. https://doi.org/10.1093/oxfordhb/9780195374148.013.0243

Friese, U., Daume, J., Göschl, F., König, P., Wang, P., & Engel, A. K. (2016). Oscillatory brain activity during multisensory attention reflects activation, disinhibition, and cognitive control. Scientific Reports, 6(1), 32775. https://doi.org/10.1038/srep32775

Ganesan, K., Plass, J., Beltz, A. M., Liu, Z., Grabowecky, M., Suzuki, S., Stacey, W. C., Wasade, V. S., Towle, V. L., Tao, J. X., Wu, S., Issa, N. P., & Brang, D. (2021). Visual speech differentially modulates beta, theta, and high gamma bands in auditory cortex. European Journal of Neuroscience, 54(9), 7301–7317. https://doi.org/10.1111/ejn.15482

Getzmann, S., Klatt, L.-I., Schneider, D., Begau, A., & Wascher, E. (2020). EEG correlates of spatial shifts of attention in a dynamic multi-talker speech perception scenario in younger and older adults. Hearing Research, 398, 108077. https://doi.org/10.1016/j.heares.2020.108077

Gevins, A. (1997). High-resolution EEG mapping of cortical activation related to working memory: effects of task difficulty, type of processing, and practice. Cerebral Cortex, 7(4), 374–385. https://doi.org/10.1093/cercor/7.4.374

Grandchamp, R., & Delorme, A. (2011). Single-Trial Normalization for Event-Related Spectral Decomposition Reduces Sensitivity to Noisy Trials. Frontiers in Psychology, 2, 1–14. https://doi.org/10.3389/fpsyg.2011.00236

Gratton, G. (2018). Brain reflections: A circuit-based framework for understanding information processing and cognitive control. Psychophysiology, 55(3), 1–26. https://doi.org/10.1111/psyp.13038

Guerreiro, M. J. S., Murphy, D. R., & Van Gerven, P. W. M. (2010). The role of sensory modality in age-related distraction: A critical review and a renewed view. Psychological Bulletin, 136(6), 975–1022. https://doi.org/10.1037/a0020731

Habak, C., Seghier, M. L., Brûlé, J., Fahim, M. A., & Monchi, O. (2019). Age Affects How Task Difficulty and Complexity Modulate Perceptual Decision-Making. Frontiers in Aging Neuroscience, 11, 1–10. https://doi.org/10.3389/fnagi.2019.00028

Jensen, O., & Tesche, C. D. (2002). Frontal theta activity in humans increases with memory load in a working memory task. European Journal of Neuroscience, 15(8), 1395–1399. https://doi.org/10.1046/j.1460-9568.2002.01975.x

Keil, J., Müller, N., Ihssen, N., & Weisz, N. (2012). On the variability of the McGurk effect: Audiovisual integration depends on prestimulus brain states. Cerebral Cortex, 22(1), 221–231. https://doi.org/10.1093/cercor/bhr125

Keil, J., & Senkowski, D. (2018). Neural Oscillations Orchestrate Multisensory Processing. The Neuroscientist, 24(6), 609–626. https://doi.org/10.1177/1073858418755352

Keller, A. S., Payne, L., & Sekuler, R. (2017). Characterizing the roles of alpha and theta oscillations in multisensory attention. Neuropsychologia, 99, 48–63. https://doi.org/10.1016/j.neuropsychologia.2017.02.021

Klatt, L.-I., Getzmann, S., Begau, A., & Schneider, D. (2020). A dual mechanism underlying retroactive shifts of auditory spatial attention: dissociating target-and distractor-related modulations of alpha lateralization. Scientific Reports, 10(1), 13860. https://doi.org/10.1038/s41598-020-70004-2

Klucharev, V., Möttönen, R., & Sams, M. (2003). Electrophysiological indicators of phonetic and non-phonetic multisensory interactions during audiovisual speech perception. Cognitive Brain Research, 18(1), 65–75. https://doi.org/10.1016/j.cogbrainres.2003.09.004

Lakens, D. (2013). Calculating and reporting effect sizes to facilitate cumulative science: a practical primer for t-tests and ANOVAs. Frontiers in Psychology, 4, 1–12. https://doi.org/10.3389/fpsyg.2013.00863

Lange, J., Christian, N., & Schnitzler, A. (2013). Audio–visual congruency alters power and coherence of oscillatory activity within and between cortical areas. NeuroImage, 79, 111–120. https://doi.org/10.1016/j.neuroimage.2013.04.064

Lopez-Calderon, J., & Luck, S. J. (2014). ERPLAB: an open-source toolbox for the analysis of event-related potentials. Frontiers in Human Neuroscience, 8(213), 1–14. https://doi.org/10.3389/fnhum.2014.00213

Luck, S. J. (2014). The Mass Univariate Approach and Permutation Statistics. In A. B. Book (Ed.), An Introduction to the Event-Related Potential Technique, Second Edition (pp. 1–15). MIT Press. http://mitp-content-server.mit.edu:18180/books/content/sectbyfn?collid=books_pres_0&fn=Ch_13_Mass_Univariate_and_Permutations_0.pdf&id=8575

Makeig, S., Debener, S., Onton, J., & Delorme, A. (2004). Mining event-related brain dynamics. Trends in Cognitive Sciences, 8(5), 204–210. https://doi.org/10.1016/j.tics.2004.03.008

Maris, E., & Oostenveld, R. (2007). Nonparametric statistical testing of EEG-and MEG-data. Journal of Neuroscience Methods, 164(1), 177–190. https://doi.org/10.1016/j.jneumeth.2007.03.024

Martin, J. S., & Jerger, J. F. (2005). Some effects of aging on central auditory processing. The Journal of Rehabilitation Research and Development, 42, 25. https://doi.org/10.1682/JRRD.2004.12.0164

Maurer, U., Brem, S., Liechti, M., Maurizio, S., Michels, L., & Brandeis, D. (2015). Frontal Midline Theta Reflects Individual Task Performance in a Working Memory Task. Brain Topography, 28(1), 127–134. https://doi.org/10.1007/s10548-014-0361-y

McGurk, H., & MacDonaldJohn. (1976). Hearing lips and seeing voices. Nature, 264(5588), 746–748. https://doi.org/10.1038/264746a0

Michail, G., Senkowski, D., Niedeggen, M., & Keil, J. (2021). Memory Load Alters Perception-Related Neural Oscillations during Multisensory Integration. The Journal of Neuroscience, 41(7), 1505–1515. https://doi.org/10.1523/JNEUROSCI.1397-20.2020

Misselhorn, J., Friese, U., & Engel, A. K. (2019). Frontal and parietal alpha oscillations reflect attentional modulation of cross-modal matching. Scientific Reports, 9(1), 5030. https://doi.org/10.1038/s41598-019-41636-w

Morís Fernández, L., Torralba, M., & Soto-Faraco, S. (2018). Theta oscillations reflect conflict processing in the perception of the McGurk illusion. European Journal of Neuroscience, 48(7), 2630–2641. https://doi.org/10.1111/ejn.13804

Morís Fernández, L., Visser, M., Ventura-Campos, N., Ávila, C., & Soto-Faraco, S. (2015). Top-down attention regulates the neural expression of audiovisual integration. NeuroImage, 119, 272–285. https://doi.org/10.1016/j.neuroimage.2015.06.052

Noguchi, K., Gel, Y. R., Brunner, E., & Konietschke, F. (2012). nparLD : An R Software Package for the Nonparametric Analysis of Longitudinal Data in Factorial Experiments. Journal of Statistical Software, 50(12), 1–23. https://doi.org/10.18637/jss.v050.i12

O’Sullivan, A. E., Lim, C. Y., & Lalor, E. C. (2019). Look at me when I’m talking to you: Selective attention at a multisensory cocktail party can be decoded using stimulus reconstruction and alpha power modulations. European Journal of Neuroscience, 50(8), 3282–3295. https://doi.org/10.1111/ejn.14425

Olusanya, B. O., Davis, A. C., & Hoffman, H. J. (2019). Hearing loss grades and the International classification of functioning, disability and health. Bulletin of the World Health Organization, 97(10), 725–728. https://doi.org/10.2471/BLT.19.230367

Owsley, C. (2011). Aging and vision. Vision Research, 51(13), 1610–1622. https://doi.org/10.1016/j.visres.2010.10.020

Passow, S., Westerhausen, R., Wartenburger, I., Hugdahl, K., Heekeren, H. R., Lindenberger, U., & Li, S.-C. (2012). Human aging compromises attentional control of auditory perception. Psychology and Aging, 27(1), 99–105. https://doi.org/10.1037/a0025667

Peelle, J. E., & Sommers, M. S. (2015). Prediction and constraint in audiovisual speech perception. Cortex, 68, 169–181. https://doi.org/10.1016/j.cortex.2015.03.006

Peelle, J. E., & Wingfield, A. (2016). The Neural Consequences of Age-Related Hearing Loss. Trends in Neurosciences, 39(7), 486–497. https://doi.org/10.1016/j.tins.2016.05.001

Pion-Tonachini, L., Kreutz-Delgado, K., & Makeig, S. (2019). ICLabel: An automated electroencephalographic independent component classifier, dataset, and website. NeuroImage, 198, 181–197. https://doi.org/10.1016/j.neuroimage.2019.05.026

Rey-Mermet, A., & Gade, M. (2018). Inhibition in aging: What is preserved? What declines? A meta-analysis. Psychonomic Bulletin & Review, 25(5), 1695–1716. https://doi.org/10.3758/s13423-017-1384-7

Roa Romero, Y., Keil, J., Balz, J., Niedeggen, M., Gallinat, J., & Senkowski, D. (2016). Alpha-Band Oscillations Reflect Altered Multisensory Processing of the McGurk Illusion in Schizophrenia. Frontiers in Human Neuroscience, 10, 1–12. https://doi.org/10.3389/fnhum.2016.00041

Roa Romero, Y., Senkowski, D., & Keil, J. (2015). Early and late beta-band power reflect audiovisual perception in the McGurk illusion. Journal of Neurophysiology, 113(7), 2342–2350. https://doi.org/10.1152/jn.00783.2014

Salthouse, T. A. (1979). Adult age and the speed-accuracy trade-off. Ergonomics, 22(7), 811–821. https://doi.org/10.1080/00140137908924659

Salthouse, T. A. (2000). Aging and measures of processing speed. Biological Psychology, 54(1–3), 35–54. https://doi.org/10.1016/S0301-0511(00)00052-1

Schepers, I. M., Schneider, T. R., Hipp, J. F., Engel, A. K., & Senkowski, D. (2013). Noise alters beta-band activity in superior temporal cortex during audiovisual speech processing. NeuroImage, 70, 101–112. https://doi.org/10.1016/j.neuroimage.2012.11.066

Schneider, D., Herbst, S. K., Klatt, L., & Wöstmann, M. (2021). Target enhancement or distractor suppression? Functionally distinct alpha oscillations form the basis of attention. European Journal of Neuroscience, 1–10. https://doi.org/10.1111/ejn.15309

Schneider, D., Zickerick, B., Thönes, S., & Wascher, E. (2020). Encoding, storage, and response preparation—Distinct EEG correlates of stimulus and action representations in working memory. Psychophysiology, 57(6), 1–15. https://doi.org/10.1111/psyp.13577

Schwartz, J.-L., & Savariaux, C. (2014). No, There Is No 150 ms Lead of Visual Speech on Auditory Speech, but a Range of Audiovisual Asynchronies Varying from Small Audio Lead to Large Audio Lag. PLoS Computational Biology, 10(7). https://doi.org/10.1371/journal.pcbi.1003743

Sekiyama, K., Soshi, T., & Sakamoto, S. (2014). Enhanced audiovisual integration with aging in speech perception: a heightened McGurk effect in older adults. Frontiers in Psychology, 5(323), 1–12. https://doi.org/10.3389/fpsyg.2014.00323

Shapiro, S. S., & Wilk, M. B. (1965). An Analysis of Variance Test for Normality (Complete Samples). Biometrika, 52(3/4), 591. https://doi.org/10.2307/2333709

Shatzer, H., Shen, S., Kerlin, J. R., Pitt, M. A., & Shahin, A. J. (2018). Neurophysiology underlying influence of stimulus reliability on audiovisual integration. European Journal of Neuroscience, 48(8), 2836–2848. https://doi.org/10.1111/ejn.13843

Simon, D. M., & Wallace, M. T. (2018). Integration and Temporal Processing of Asynchronous Audiovisual Speech. Journal of Cognitive Neuroscience, 30(3), 319–337. https://doi.org/10.1162/jocn_a_01205

Sommers, M. S., Tye-Murray, N., & Spehar, B. (2005). Auditory-Visual Speech Perception and Auditory-Visual Enhancement in Normal-Hearing Younger and Older Adults. Ear and Hearing, 26(3), 263–275. https://doi.org/10.1097/00003446-200506000-00003

Stekelenburg, J. J., & Vroomen, J. (2007). Neural Correlates of Multisensory Integration of Ecologically Valid Audiovisual Events. Journal of Cognitive Neuroscience, 19(12), 1964–1973. https://doi.org/10.1162/jocn.2007.19.12.1964

Stothart, G., & Kazanina, N. (2016). Auditory perception in the aging brain: the role of inhibition and facilitation in early processing. Neurobiology of Aging, 47, 23–34. https://doi.org/10.1016/j.neurobiolaging.2016.06.022

van Wassenhove, V., Grant, K. W., & Poeppel, D. (2005). Visual speech speeds up the neural processing of auditory speech. Proceedings of the National Academy of Sciences, 102(4), 1181–1186. https://doi.org/10.1073/pnas.0408949102

Weiss, S., & Mueller, H. M. (2012). “Too Many betas do not Spoil the Broth”: The Role of Beta Brain Oscillations in Language Processing. Frontiers in Psychology, 3(JUN), 1–15. https://doi.org/10.3389/fpsyg.2012.00201

Weisz, N., Hartmann, T., Müller, N., Lorenz, I., & Obleser, J. (2011). Alpha Rhythms in Audition: Cognitive and Clinical Perspectives. Frontiers in Psychology, 2(APR), 1–15. https://doi.org/10.3389/fpsyg.2011.00073

Wilcoxon, F. (1946). Individual Comparisons of Grouped Data by Ranking Methods. Journal of Economic Entomology, 39(2), 269–270. https://doi.org/10.1093/jee/39.2.269

Winneke, A. H., & Phillips, N. A. (2011). Does audiovisual speech offer a fountain of youth for old ears? An event-related brain potential study of age differences in audiovisual speech perception. Psychology and Aging, 26(2), 427–438. https://doi.org/10.1037/a0021683

Wong, P. C. M., Ettlinger, M., Sheppard, J. P., Gunasekera, G. M., & Dhar, S. (2010). Neuroanatomical Characteristics and Speech Perception in Noise in Older Adults. Ear & Hearing, 31(4), 471–479. https://doi.org/10.1097/AUD.0b013e3181d709c2

Wöstmann, M., Herrmann, B., Maess, B., & Obleser, J. (2016). Spatiotemporal dynamics of auditory attention synchronize with speech. Proceedings of the National Academy of Sciences, 113(14), 3873–3878. https://doi.org/10.1073/pnas.1523357113

Wostmann, M., Herrmann, B., Wilsch, A., & Obleser, J. (2015). Neural Alpha Dynamics in Younger and Older Listeners Reflect Acoustic Challenges and Predictive Benefits. Journal of Neuroscience, 35(4), 1458–1467. https://doi.org/10.1523/JNEUROSCI.3250-14.2015

Zion Golumbic, E., Cogan, G. B., Schroeder, C. E., & Poeppel, D. (2013). Visual Input Enhances Selective Speech Envelope Tracking in Auditory Cortex at a “Cocktail Party.” Journal of Neuroscience, 33(4), 1417–1426. https://doi.org/10.1523/JNEUROSCI.3675-12.2013

